# Carbon pool trends and dynamics within a subtropical peatland during long-term restoration

**DOI:** 10.1101/196907

**Authors:** Paul Julian, Stefan Gerber, Alan L. Wright, Binhe Gu, Todd Z. Osborne

## Abstract

**Background:** The Florida Everglades has undergone significant ecological change spanning the continuum of disturbance to restoration. While the restoration effort is not complete and the ecosystem continues to experience short duration perturbations, a better understanding of long-term C dynamics of the Everglades is needed to facilitate new restoration efforts. This study evaluated temporal trends of different aquatic carbon (C) pools of the northern Everglades Protection Area over a 20-year period to gauge historic C cycling patterns. Dissolved inorganic C (DIC), dissolved organic C (DOC), particulate organic C (POC), and surface water carbon dioxide (pCO_2(aq)_) were investigated between May 1, 1994 and April 30, 2015.

**Results:** Annual mean concentrations of DIC, DOC, POC, and pCO_2(aq)_ significantly decreased through time or remained constant across the Water Conservation Areas (WCAs). Overall, the magnitude of the different C pools in the water column significantly differed between regions. Outgassing of CO_2_ was dynamic across the Everglades ranging from 420 to 2001 kg CO_2_ yr-1. Overall the historic trend in CO_2_ flux from the marsh declined across our study area while pCO_2(aq)_ largely remained somewhat constant with the exception of Water Conservation Area 2 which experienced significant declines in pCO_2(aq)_. Particulate OC concentrations were consistent between WCAs, but a significantly decreasing trend in annual POC concentrations were observed.

**Conclusions:** Hydrologic condition and nutrient inputs significantly influenced the balance, speciation, and flux of C pools across WCAs suggesting a subsidy-stress response in C dynamics relative to landscape scale responses in nutrient availability. The interplay between nutrient inputs and hydrologic condition exert a driving force on the balance between DIC and DOC production via the metabolism of organic matter which forms the base of the aquatic foodweb. Along the restoration trajectory as water quality and hydrology continues to improve it is expected that C pools will respond accordingly.

## Introduction

Wetlands, including peatlands, are net C sinks that store a large amount of the global C created by an unbalanced accumulation of C via plant productivity and export from decomposition of organic matter via C dioxide (CO_2_) or methane (CH_4_) to the atmosphere. Carbon forms such as dissolved inorganic C (DIC), particulate organic C (POC) and DOC can be transported laterally through run-off (Updegraff et al. 1995; Freeman et al. 2004; Billett and Moore 2008), and contribute to wetlands’ carbon budgets. This type of export C, more specifically organic species from wetlands, represents a significant regional redistribution of terrestrial C and exerts important controls on aquatic productivity in downstream waterbodies. Dissolved organic and inorganic C may thereby enter a feedback loop with aquatic productivity within a wetland (Carpenter and Pace 1997; Hanson et al. 2003).

Net aquatic production (NAP), or total metabolic balance of an aquatic ecosystem, is the difference between gross primary production and ecosystem respiration in the water column. As a result of maintaining a metabolic balance, different “pools” of aquatic carbon are influenced by ecosystem processes related to ecosystem metabolism and function. Aquatic ecosystems with low nutrients such as total phosphorus (TP) and high DOC from either allochthonous (external) or autochthonous (internal) sources tend to be net heterotrophic, while those exhibiting low DOC and high TP tend towards net autotrophy (Hanson et al. 2003; Waiser and Robarts 2004). Therefore, the supply of various C pools and other water quality indicators are important in understanding ecosystem function.

Most wetlands and lake ecosystems are heterotrophic with respect to aquatic metabolism with surface waters acting as net sources of CO_2_ to the atmosphere by decomposition of organic matter (Kling et al. 1991; Cole et al. 1994; Waiser and Robarts 2004). In contrast, some coastal wetlands and a small number of lakes are considered autotrophic (Kling et al. 1992; Cole et al. 1994; Duarte and Cebrian 1996; Gu et al. 2011; Hopkinson et al. 2012). The net C source in wetland and littoral zones could be sustained through C fixation by emergent vegetation and its subsequent decomposition. Thus, while the surface water may be heterotrophic, the wetland ecosystem as a whole (water + soil), can still function as a net C sinks resulting in overall C accretion.

The major sources of total dissolved CO_2_ contributing to aquatic systems are (1) diffusion of atmospheric CO_2_ from the atmosphere, (2) hydrologic transport from upstream, (3) groundwater discharge, (4) CO_2_ derived from organic matter decomposition within the water column and the soil and (5) the dissolution of C minerals (Telmer and Veizer 1999; Finlay 2003). In some river and small stream systems, metabolism of C inputs appears to be greater than *in-situ* primary production resulting in higher aqueous dissolved CO_2_ than CO_2_ in the atmosphere (Cole et al. 1994). A total metabolic balance is driven by factors that govern both primary production and respiration. Productivity can be influenced by nutrients, light availability, thermal cycle, hydrodynamics (i.e. current, velocity, mixing, etc.), internal loading of organic matter and food web structure. Meanwhile ecosystem respiration may be strongly subsidized by external loading of organic matter and nutrients (Odum 1956; Kling et al. 1991; Wetzel 1992; Cole et al. 2000; Hessen et al. 2004; Elser et al. 2007; Gu et al. 2011). Therefore, the combination and interaction of these factors influence the partitioning and conversion of C into distinct pools.

The objectives of this study were to evaluate different aquatic C pools within the Everglades ecosystem. The first objective of this study was to evaluate long-term regional trends in aquatic C (DIC, DOC, POC and *p*CO_2_) within the Everglades as large scale restoration efforts progress to restore quality, quantity, and timing of water to the Everglades ecosystem. The first hypothesis is as the system recovers from degraded water quality impacts, C pools will decline or level out. The decrease may be directly caused by reduced nutrient availability to primary producers and possibly modified by altered (increasing or decreasing) organic matter turnover in the water column and soils. The second objective is to estimate CO_2_ flux from the Everglades Protection Area (EPA) marsh to the atmosphere and investigate long-term trends, with the hypotheses that aquatic CO_2_ fluxes will also decline concurrently with the decline in other C pools as a response to restoration.

## Methods

### Study Area

The Everglades ecosystem is a complex system of marshes, canals and levees with water control structures covering approximately 10,000 km^2^ of former contiguous Everglades marsh and currently divided into large, distinct shallow impoundments (Bancroft et al. 1992; Light and Dineen 1994). Surface water is delivered primarily from the north and west to the EPA through water control structures connecting the Everglades Agricultural Area (EAA), stormwater treatment areas (STAs) and the westernbasins and through urban areas along the eastern edge of the EPA. Surface water from these land areas are delivered to the EPA through water control structures, but the timing and distribution of the surface water inflows from the upstream watersheds to the EPA are based on complex series of operational decisions accounting for natural and environmental system requirements, water supply for urbanized and natural areas, aquifer recharge, and flood control (Julian et al. 2016).

This study was centered on the northern third of the EPA, known as the water conservation areas (WCAs; Fig. 1) that receive the majority of surface water inputs through the highly managed surface water system via a combination of canals and water control structures. Water Conservation Area 1 is unique in the distinct point source inputs of surface water that moves around the marsh edge via the perimeter canal. As a result of this limited interaction of the WCA-1 marsh interior with mineral-rich canal drainage water, WCA-1 is the sole remaining soft-water ecosystem in the Everglades and has a predominately rainfall driven hydrology in the marsh interior (Newman and Hagerthey 2011). Both WCA-2 and WCA-3 surface hydrology is controlled by a system of levees and water control structures along the perimeter. Additionally canals, levees and water control structures bisect WCA-3 effectively dividing the area into four hydrologically distinct areas (DeBusk et al. 2001; Bruland et al. 2006). Soil within the WCAs are Histosols, and encompass both Loxahatchee and Everglades peat formations with depths ranging from ~1 to 2 m with a mosaic of aquatic sloughs, expanses of wet prairie, strands of sawgrass (*Cladium jamaicense* Crantz), and patches of brush and tree islands (Gleason et al. 1974; Brandt et al. 2000; DeBusk et al. 2001; Bruland et al. 2006)

**Fig. 1.**
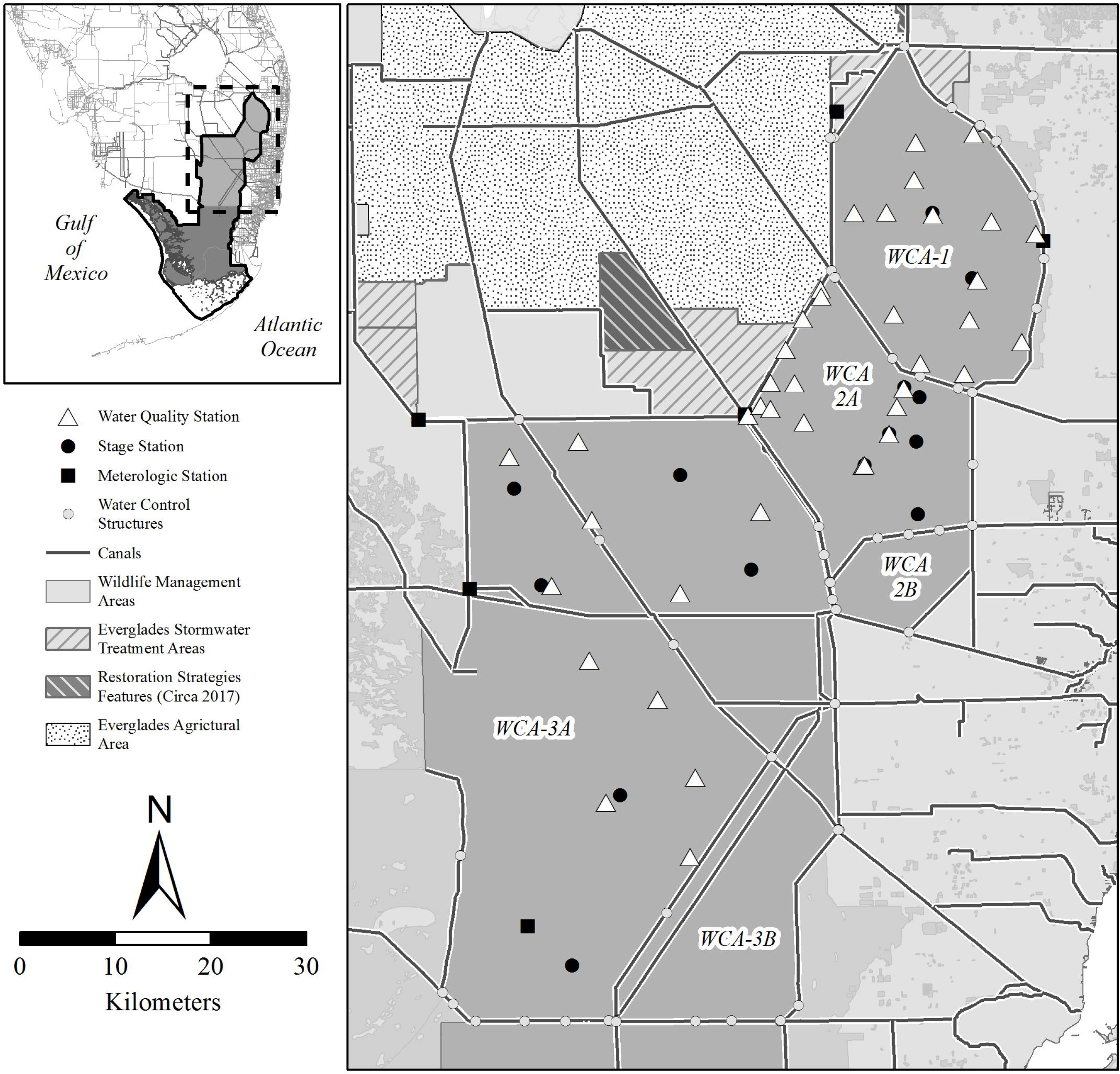
Map of surface water quality, stage and weather monitoring location within the Everglades Protection Area relative to surrounding features including the Everglades Stormwater Treatment Area, Everglades Agricultural Area and Water Conservation Areas.

### Source of Data

Water quality data was retrieved from the South Florida Water Management District (SFWMD) online database (DBHYDRO; www.sfwmd.gov/dbhydro) for sites within the WCAs identified in Fig. 1 between water year 1995 to 2015 (May 1, 1994 – April 30, 2015). Only stations with greater than four samples per year and three years of monitoring data were considered for this analysis resulting in 44 monitoring stations across the EPA. Surface water grab samples were typically collected between 06:00 to 17:00 local time, with most samples collected at approximately 09:00. Water quality parameters used include alkalinity, pH, temperature, specific conductivity, DOC, total phosphorus (TP) and total nitrogen (TN). In order to quantify air-water net CO_2_ exchange between surface waters and the atmosphere additional data on wind speed and atmospheric CO_2_ partial pressure was obtained. Wind speed data measured 10 meters above ground elevation was retrieved from metrological stations within the EPA identified by Fig. 1. Monthly atmospheric partial pressure carbon dioxide (*p*CO_2(atm)_) data was retrieved from the National Oceanic & Atmospheric Administration (NOAA) online global monitoring division webpage (www.esrl.noaa.gov/gmd) using the Key Biscayne monitoring location (25.6654° N, 80.1580° W; NAD1983). Stage elevation data was retrieved from the SFWMD DBHYDRO for stage monitoring site identified in Fig. 1 between water year (WY) 1979 to 2015 (May 1, 1978 – April 30, 2015).

Water quality data were screened based on laboratory qualifier codes, consistent with FDEP’s quality assurance rule (Florida Administrative Code 2008). Any datum associated with a fatal qualifier indicating a potential data quality problem were removed from the analysis. For purposes of data analysis and summary statistics, data reported as less than method detection limit (MDL) were assigned a value of one-half the MDL, unless otherwise noted.

### Data Analysis

Dissolved inorganic C concentrations were calculated from the relationship between water column alkalinity, pH and temperature as outlined by Wetzel and Likens (2000). This methodology of quantifying DIC concentrations is consistent with previous studies where they determined that the calculated DIC concentration based on alkalinity and relevant parameters can be used to estimate DIC concentration in the absence of direct measure DIC concentration (Gu et al. 2008, 2011). Particulate organic C concentration were calculated from the difference between total organic C (TOC) and DOC. The concentration of dissolved free CO_2_ within the water column were calculated using the pH and CO_2_ fraction relationship presented in (Wetzel and Likens 2000) (Table 1, equation 1). Surface water *p*CO_2_ was calculated using Henry’s Law (Table 1, equation 2) where K_H_ is the dissolution constant of CO_2_ corrected for water temperature (Table 1, equation 3). Atmospheric concentration of CO_2_ [CO_2(atm)_] above the stagnant layer were estimated from Henry’s Law using monthly atmospheric CO_2_ partial pressure from the Key Biscayne NOAA monitoring location (Table 1, equation 4). Due to the limited availability of wind data CO_2_ flux calculations were limited to a nine-year period (WY1999 - 2008). During the nine-year period the WCAs experienced changes in climate (i.e. drought and flood), water quality and system operations due to the construction and operation of the Everglades Stormwater Treatment Areas. Therefore, to extend the period of record of flux calculations comparisons of calculations with differences in wind speed data were performed (Appendix 1). As a result of this comparison, a period of record mean wind speed value of 2.87 m s^-1^ was substituted for wind speed in the flux calculations to estimate the flux of CO_2_ between the atmosphere and surface water, equation 5 (Table 1) was used which incorporated CO_2_ diffusion coefficient (Table 1, equation 6) and surface boundary layer thickness (Table 1, equation 7). For sites with dense macrophyte coverage, it was assumed that the emergent macrophytes would reduce wind speed at the air-water interface to effectively zero similar to (Hagerthey et al. 2010). Water column DIC concentrations and surface water *p*CO_2_ (*p*CO_2(aq)_) were calculated for all station with the necessary data between WY1995 to WY2015.

**Table 1.**
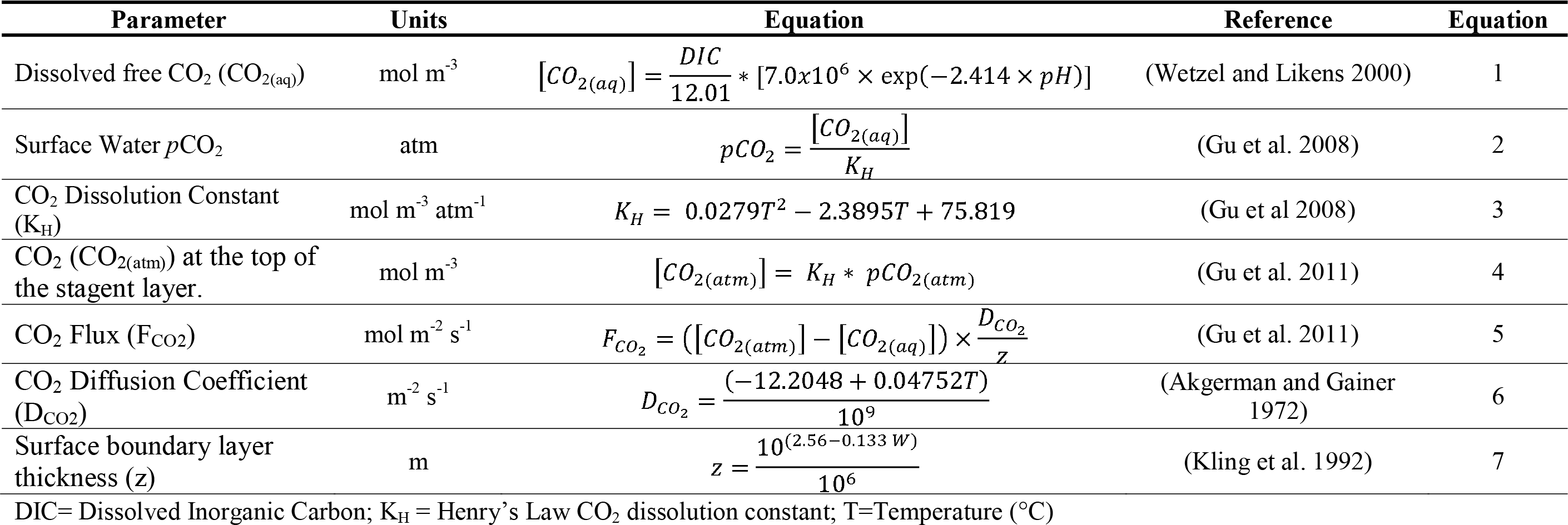
Equations used to calculate surface water *p*CO_2_ concentrations and CO_2_ flux rates.

Regional comparison was conducted using annual mean DIC, DOC, POC and *p*CO_2(aq)_ concentrations for each WCA based on the Florida water year (May-April). Station specific trend analysis was using Kendall’s τ correlation analysis and Thiel-Sen’s slope estimate (zyp R-package; (Bronaugh and Werner 2013). Annual mean DIC, DOC, POC, *p*CO_2(aq)_ and CO_2_ flux were compared between WCAs using the Kruskal-Wallis rank sum test and Dunn’s test of multiple comparisons (Dunn’s test R-package) (Dinno 2015) using stations with greater than four samples collected in a given WY. Annual mean DOC and DIC ratio were computed using mass per volume concentrations and compared between WCAs using the Kruskal-Wallis rank sum test and Dunn’s test of multiple comparisons. Unless otherwise noted all statistical operations were performed using the base stats R-package. All statistical operations were performed with R© (Ver 3.1.2, R Foundation for Statistical Computing, Vienna Austria) at a critical level of significance of α = 0.05.

## Results

During this study, calculated DIC concentrations ranged from 1.9 mg C L^−1^ to 144.0 mg C L^-1^ across the WCAs. Statistically significantly declining annual DIC trends were observed at a total of 9 stations during this study, no trend was apparent for 32 of the stations during this study, the remaining didn’t have sufficient data to conduct a trend analysis (Fig 2). Most notable significantly decreasing trends were observed at stations along the primary eutrophication gradient within WCA-2 where historically untreated stormwater runoff was discharged resulting in a significant area of impact (DeBusk et al. 2001). Regional annual mean DIC concentrations between WY1995 and 2015 significantly declined in WCA-2 at a rate of – 0.53 mg L^-1^ yr^-1^, no temporal trend was apparent in WCA-1 and WCA-3 (Table 2). Dissolved inorganic C concentration significantly differed between WCAs (χ^2^=54.8, df=2, ρ<0.001) with DIC concentration being lowest within WCA-1 (17.2 ± 0.6 mg C L^-1^) and greatest in WCA-2 (67.1 ± 0.7 mg C L^-1^) (Fig. 3). Monthly mean DIC and stage elevation was significantly negatively correlated for WCA-3; negatively correlated with mean monthly water temperature for WCA-2 and WCA-3; positively correlated with mean monthly TP concentration for WCA-1 and WCA-2; TN and specific conductance for all regions of the study area (Table 4).

**Fig. 2.**
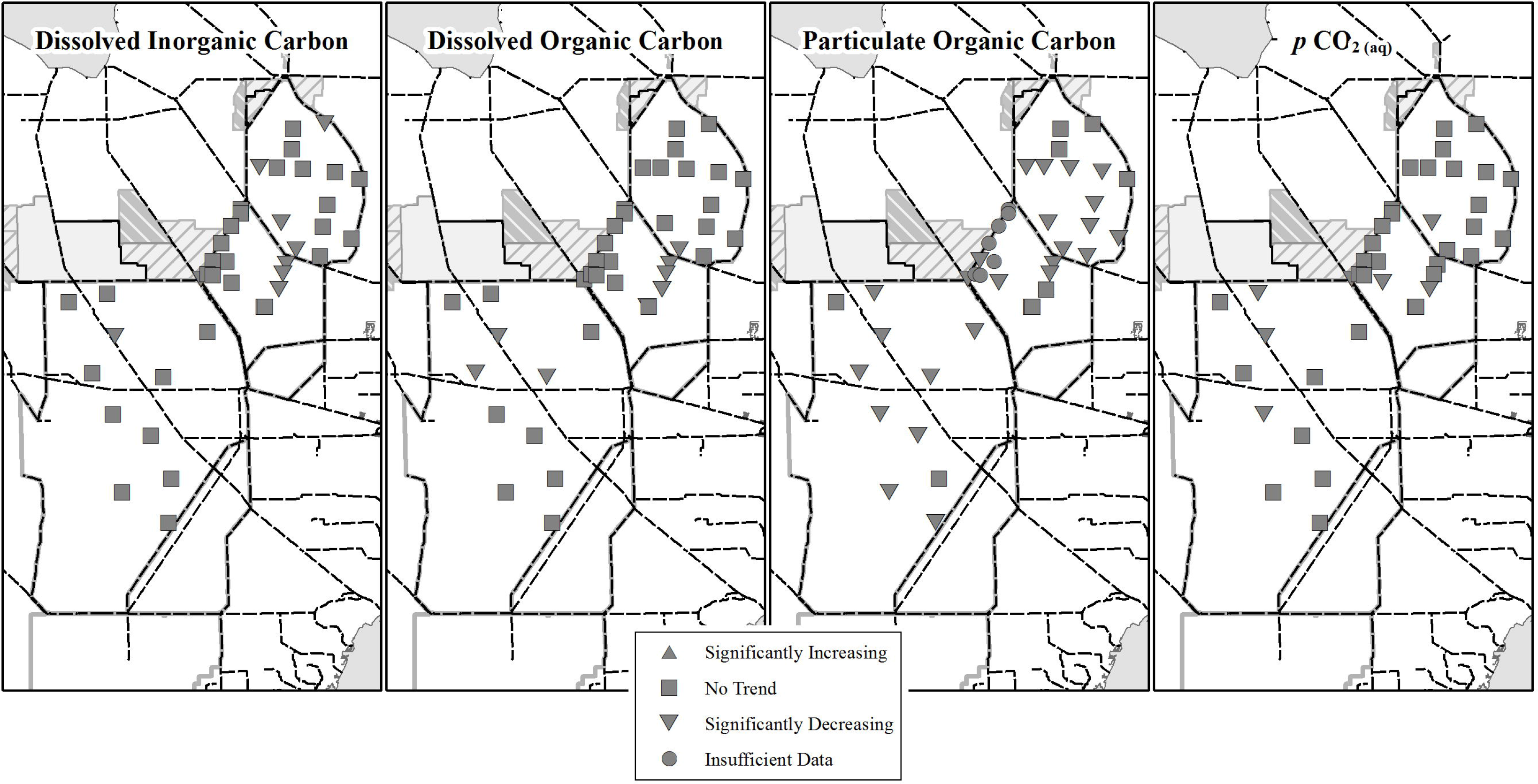
Annual Kendall trend analysis results for each monitoring location with enough data (greater than six samples per year and three water years) for dissolved inorganic carbon, dissolved organic carbon, particulate organic carbon and surface water *p*CO_2_ (*p*CO_2(aq)_). Significantly increasing or decreasing trends were determined based on a ρ-value <0.05 and positive or negative Kendall τ values to denote direction. No trend results were identified as ρ-values>0.05.

**Table 2.**
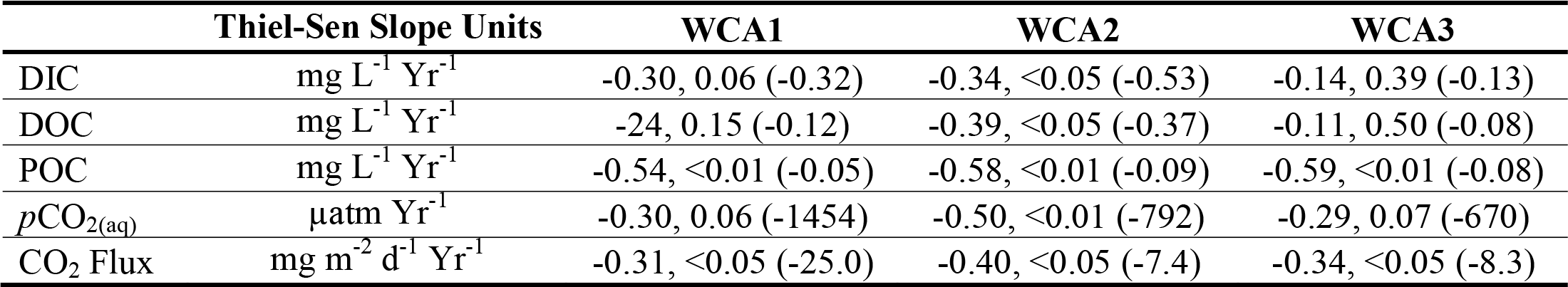
Trend analysis results expressed as Kendall’s τ, ρ-value (Thiel-Sen Slope) for annual mean dissolved inorganic carbon (DIC), dissolved organic carbon (DOC), particulate organic carbon (POC), surface water *p*CO_2_ (*p*CO_2(aq)_) and CO_2_ flux rate for each region of the Everglades Protection Area between water years 1995 and 2015 (May 1, 1994 – April 30, 2015).

**Fig. 3.**
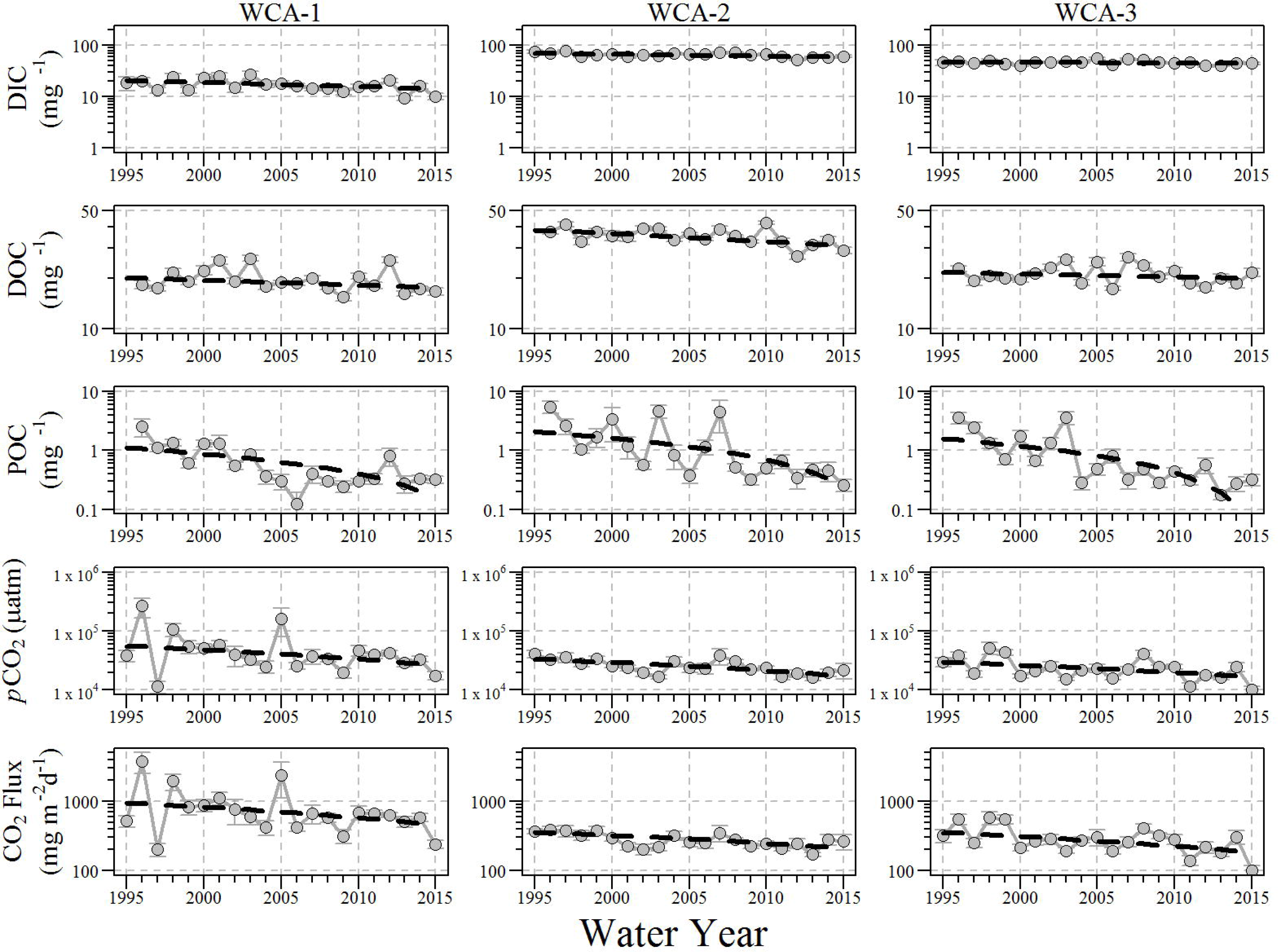
Annual mean dissolved inorganic carbon concentrations, dissolved organic carbon, particulate organic carbon and surface water *p*CO_2(aq)_ for interior portions of the Everglades Protection Area between water years 1995 and 2015 (May 1, 1994 – April 30, 2015).

Dissolved organic C concentrations ranged from 0.5 mg C L^-1^ to 65.9 mg C L^-1^ across WCAs. Significantly declining trend were observed at the individual site scale, with a total of 9 sites with significantly declining trends, meanwhile no trend was apparent for 33 of the stations in this study (Fig. 2). Regional annual mean DOC concentrations significantly declined in WCA-2 at a rate of – 0.37 mg L^-1^ yr^-1^, no temporal trend was apparent in WCA-1 and WCA-3 (Table 2). Annual mean DOC concentrations significantly differed between WCAs (Fig. 4, χ^2^=41.1 df=2, ρ<0.01), with WCA-1 and WCA-3 regional concentrations being similar to each other (19.4 ± 0.2 mg C L^-1^ and 22.1 ± 0.2 mg C L^-1^, respectively) and WCA-2 being different between WCA-1 and WCA-3 with a higher observed regional average DOC concentration (35.8 ± 0.3 mg C L^-1^) (Fig. 3). Monthly mean DOC concentrations were negatively correlated with mean monthly stage elevations for WCA-2 and WCA-3; no correlated with mean monthly surface water temperature; positively correlated with TP for WCA-2; and positively correlated with TN and specific conductivity for all regions (Table 4). Annual mean DOC:DIC values significantly differed between WCAs (Fig. 5, χ^2^=52.8, df=2, ρ<0.01) with WCA-1 having the highest observed ratio of 1.95 ± 0.07 followed by WCA-2 (0.55 ± 0.004) and WCA-3 (0.46 ± 0.003).

**Fig. 4.**
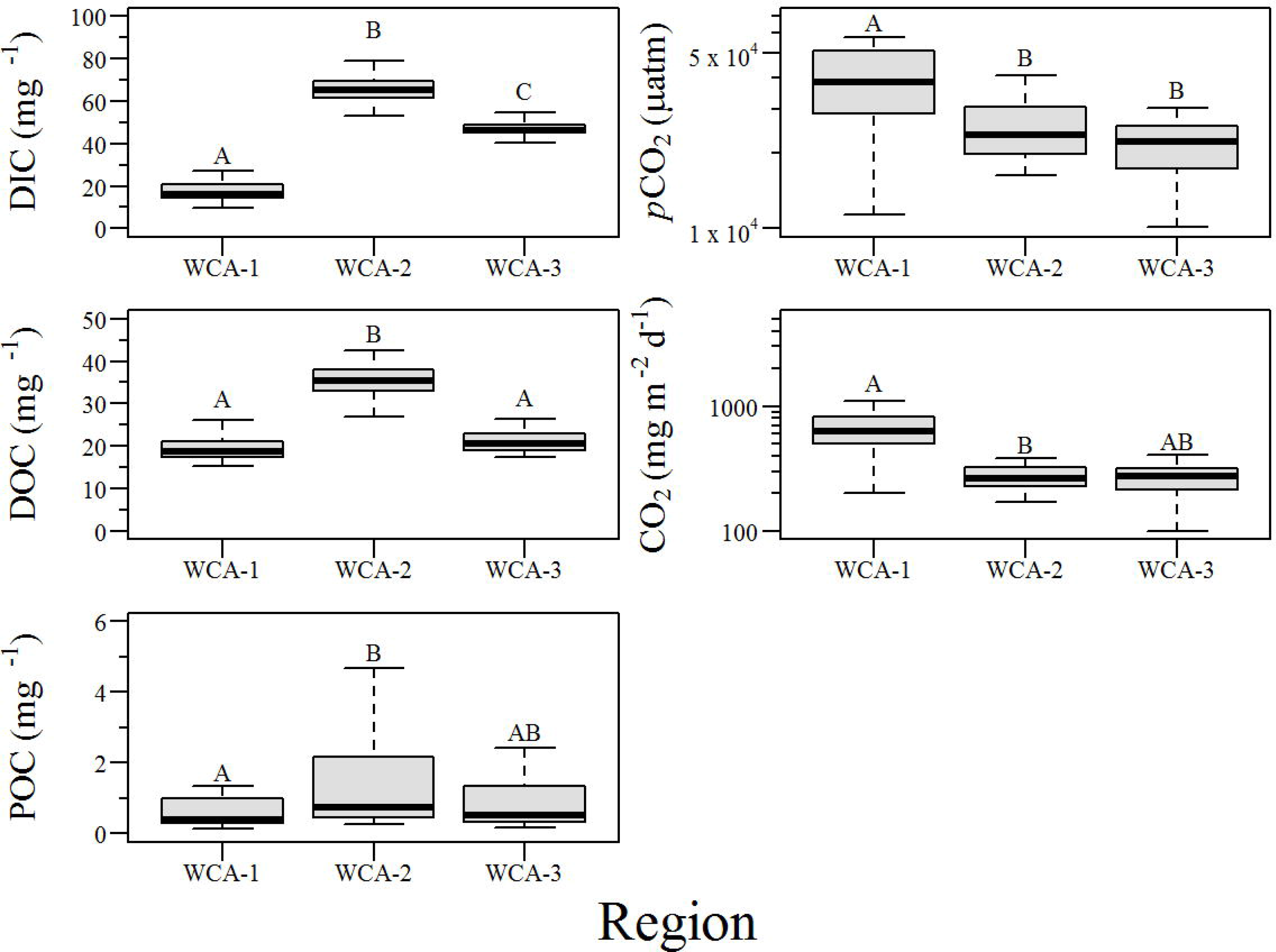
Boxplot of annual mean dissolved inorganic concentration, dissolved organic carbon, particulate organic carbon and surface water *p*CO_2(aq)_ concentration for each region of the Everglades Protection Area. Letters above box-and-whicker plots indicate statistical differences based on Dunn test results.

**Fig. 5.**
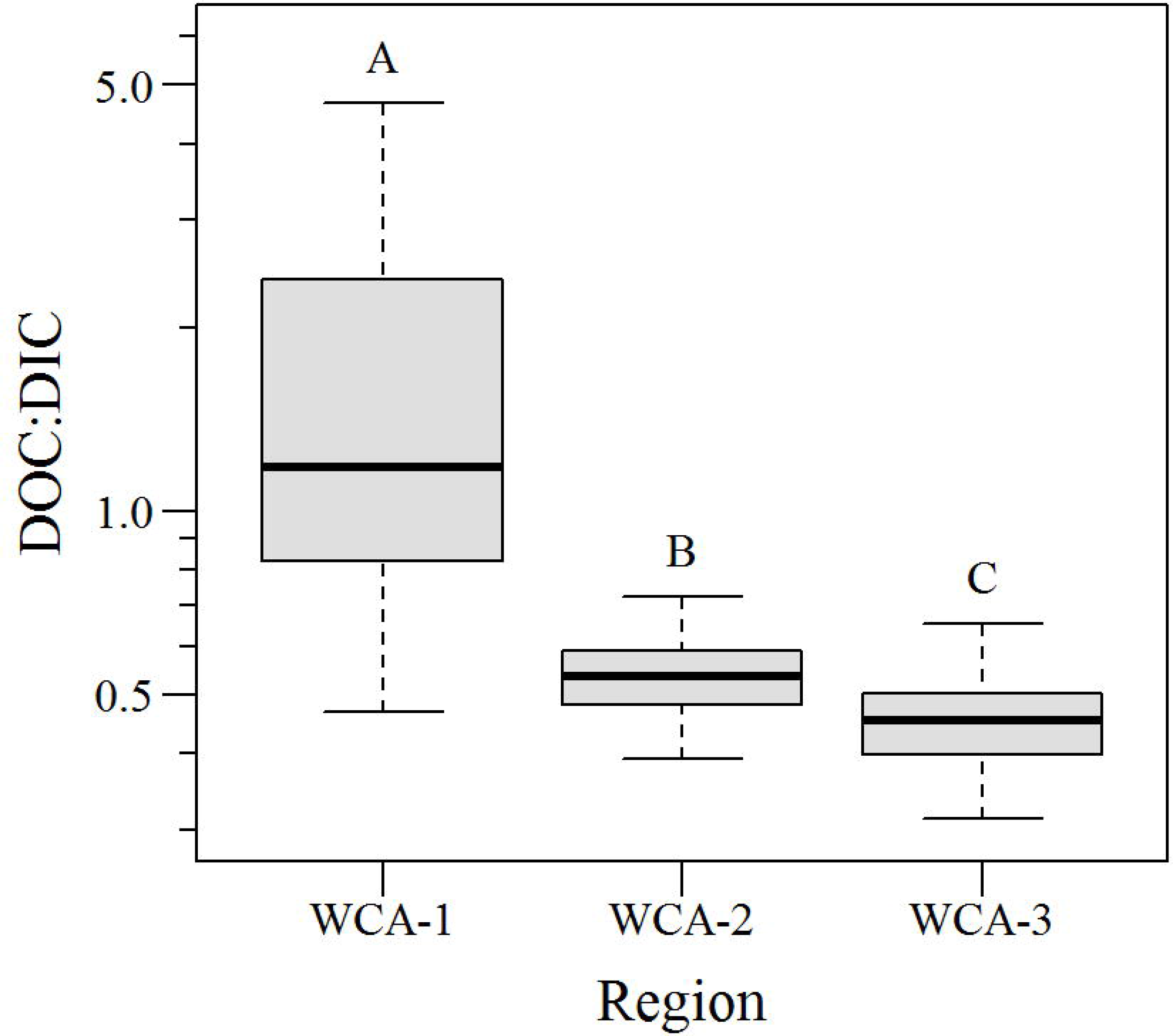
Boxplot of Dissolved Organic Carbon and Dissolved Inorganic Carbon annual mean concentration ratio for each region of the Everglades Protection Area. Graphical representation of this data can be found in Supplemental Information.

Particulate organic C concentrations ranged from 0 to 40.3 mg C L^-1^ across the WCAs. Significantly declining trends in annual mean POC concentrations were observed at 24 stations, meanwhile no trend was apparent for 9 of the stations in this study (Fig. 3). Regionally annual mean POC concentrations across all three WCA’s significantly declined at a rate of -0.26 to - 0.008 mg C L^-1^ yr^-1^ (Table 2). Annual mean POC concentrations did not significantly differ across the WCAs (Fig. 4, χ^2^=4.36, df=2, ρ=0.11), however the multiple comparison test determined a significant difference of POC concentration between WCA-1 and WCA-2 (z-score=-2.06, ρ<0.05). Regional average concentrations ranging from 0.7 ± 0.04 mg C L^-1^ (WCA-1) to 1.6 ± 0.20 mg C L^-1^ (WCA-2) with all regions experiencing significantly declining trends (Table 2). Mean monthly POC concentrations were only correlated with monthly mean TP and specific conductivity for WCA-2 and WCA-3 (Table 4).

Calculated *p*CO_2(aq)_ concentrations ranged from 135.0 μatm to 2.1 atm (2.1 x 10^6^ μatm) across the WCAs. Significantly declining trends in annual mean *p*CO_2(aq)_ were observed at 7 stations, while no trend was apparent at 34 stations during the course of this study (Fig. 2). At several stations, concurrent declines in *p*CO_2(aq)_, DIC, DOC and POC were observed potentially indicating a change in C dynamics at these specific locations within the Everglades system. Regionally annual mean *p*CO_2(aq)_ significantly declined across WCA-2 at a rate of -792 μatm yr^−1^ (Table 2) and no significant temporal trend was apparent for WCA-1 and WCA-3 (Table 2). Annual mean *p*CO_2(aq)_ significantly differed between WCAs (Fig. 4, χ^2^=13.4, df=2, ρ<0.01) with WCA-2 and WCA-3 being statistically similar (z-score=0.67, ρ=0.25; Fig. 4). Annual mean *p*CO_2(aq)_ concentrations followed a decreasing north-to-south trend with WCA-1 having the greatest concentration (52,379 ± 6,445 μatm) and WCA-3 having the lowest concentration (24,963 ± 983 μatm). Mean monthly *p*CO_2(aq)_ were positively correlated with surface water temperature within WCA-2 and positively correlated with TP for WCA-1 and WCA-2 (Table 4).

The calculated water-air CO_2_ flux ranged from -6.2 to 36,361 mg m^-2^ d^-1^ across the WCAs. Annual mean regional flux range from 285 to 876 mg m^-2^ d^-1^. Annual mean CO_2_ flux significantly declined during the course of the study for all regions (Table 2). Similar to *p*CO_2(aq)_ regional trends, annual mean flux was significantly different between WCAs (χ^2^=26.2, df=2, ρ<0.01) with annual mean flux rates being significantly different between WCA-1 and WCA-2 and WCA-3 and similar between WCA-2 and WCA-3 (z-score=0.008, ρ=0.50). Flux rates were greatest for WCA-1, followed by WCA-2 and WCA-3 (Table 3). Using the area of each region (WCA-1: 567 km^2^; WCA-2: 537 km^2^ and’ WCA-3: 2,368 km^2^) and the annual mean daily CO_2_ flux rate extrapolated to an annual estimate for each region an estimated CO_2_ flux to the atmosphere is 1,261 ± 459 kg CO_2_ yr^-1^ during this study, with the expectation that mass transport from the marsh will decreases as indicated by the annual CO_2_ flux trends.

**Table 3.**
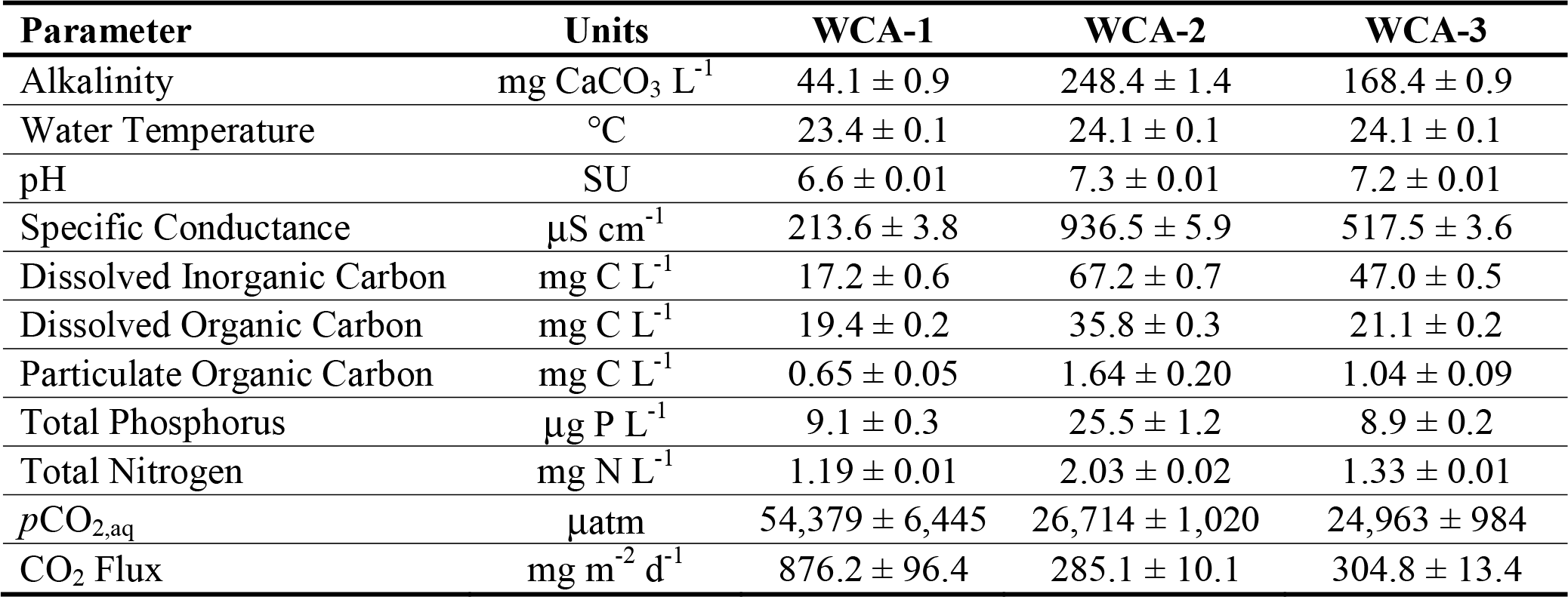
Mean ± standard error surface water parameters observed during this study within each portion of the Everglades Protection Area between water years 1995-2015 (May 1, 1994 – April 30, 2015).

| **Parameter** | **Units** | **WCA-1** | **WCA-2** | **WCA-3** |
| --- | --- | --- | --- | --- |
| Alkalinity | mg CaCO<sub>3</sub> L<sup>-1</sup> | 44.1 ± 0.9 | 248.4 ± 1.4 | 168.4 ± 0.9 |
| Water Temperature | °C | 23.4 ± 0.1 | 24.1 ± 0.1 | 24.1 ± 0.1 |
| pH | SU | 6.6 ± 0.01 | 7.3 ± 0.01 | 7.2 ± 0.01 |
| Specific Conductance | μS cm<sup>-1</sup> | 213.6 ± 3.8 | 936.5 ± 5.9 | 517.5 ± 3.6 |
| Dissolved Inorganic Carbon | mg C L<sup>-1</sup> | 17.2 ± 0.6 | 67.2 ± 0.7 | 47.0 ± 0.5 |
| Dissolved Organic Carbon | mg C L<sup>-1</sup> | 19.4 ± 0.2 | 35.8 ± 0.3 | 21.1 ± 0.2 |
| Particulate Organic Carbon | mg C L<sup>-1</sup> | 0.65 ± 0.05 | 1.64 ± 0.20 | 1.04 ± 0.09 |
| Total Phosphorus | μg P L<sup>-1</sup> | 9.1 ± 0.3 | 25.5 ± 1.2 | 8.9 ± 0.2 |
| Total Nitrogen | mg N L<sup>-1</sup> | 1.19 ± 0.01 | 2.03 ± 0.02 | 1.33 ± 0.01 |
| *p*CO<sub>2,aq</sub> | μatm | 54,379 ± 6,445 | 26,714 ± 1,020 | 24,963 ± 984 |
| CO<sub>2</sub> Flux | mg m<sup>-2</sup> d<sup>-1</sup> | 876.2 ± 96.4 | 285.1 ± 10.1 | 304.8 ± 13.4 |

Monthly *p*CO_2(atm)_ concentration ranged from 354 μatm to 404 μatm at the Key Biscayne monitoring location between WY1995 to WY2015 with a mean ± standard error of 380 ± 0.8 μatm during this period. During this period, WY mean *p*CO_2(atm)_ concentrations significantly increased (τ=1.00, ρ<0.001) during the study at a rate of 2.0 μatm year^−1^.

## Discussion

### Dissolved Inorganic Carbon and pCO_2(aq)_

Dissolved inorganic C concentrations were consistent with those reported by (Gu et al. 2008) who investigated the role of fire on C balance within WCA-2. Across the entire period of record DIC concentrations gradually decreased both at the regional (Fig. 3 and Table 2) and individual station scale (Fig. 2) with concurrent changes in water quality systems wide (Julian et al. 2016). Dissolved inorganic C is an essential source of C to benthic macrophytes and other autotrophic species (Raven et al. 1982; Sand-Jensen and Frost-Christensen 1998). Changes in water column nutrient concentrations, DIC concentrations, and water quantity influence wetland productivity which in turn influences biomass turnover, C demand in the aquatic ecosystems, and ecosystem function (Findlay et al. 2002; Bossio et al. 2006; Corstanje et al. 2007).

Each WCA has unique biogeochemical properties and hydrologic dynamics therefore DIC concentrations and C dynamics observed during this study were expected to vary between WCAs (Fig. 4). The interior portions of the WCA-1 marsh are hydrologically dominated by rainfall with very little surface water flows penetrating beyond the outer edge of the marsh (Harvey and McCormick 2009). This hydrologic setting results in low water column pH, low alkalinity and oligotrophic conditions with respect to TP (Julian et al. 2016). As such, these conditions allow for greater flux of CO_2_ from the marsh within WCA-1 relative to the other portions of the EPA presumably due to a combination of organic matter decomposition and dissolution of calcium carbonate driven by low pH conditions. Meanwhile, a different set of drivers are present for WCA-2 which historically received large quantities of storm water run-off from the EAA resulting in large areas considered to be eutrophic (DeBusk et al. 2001). However, hydrologic restorations efforts and the construction and operation of the Everglades STAs have reduced storm water inputs and TP into WCA-2 (Julian 2015). Unlike the other two WCAs, WCA-3 is hydrologically and physically compartmentalized resulting in a mosaic of areas with drastically different hydroperiods (north versus south), nutrient inputs and cycling patterns (Reddy et al. 1998; Bruland et al. 2006). The strength of the groundwater connection within WCAs is variable as is the thickness of peat/soil and depth to the lime rock bedrock across the landscape (Scheidt and Kalla 2007). As in stream ecosystems, wetland DIC is derived from several sources including the dissolution of carbonate, respiration by aquatic plants and heterotrophs by the consumption of organic matter, shallow groundwater inputs from elevated levels of soil CO_2_ and atmospheric draw-down (Wetzel 1992; Palmer et al. 2001). In some stream ecosystems it has been observed that DIC is predominately supplied and controlled by drainage of CO_2_-rich shallow groundwater into the water column (Palmer et al. 2001; Finlay 2003). In wetlands respiration by aquatic plants and heterotrophs is the dominate source and potential control of DIC concentrations (Richey et al. 2002; Hagerthey et al. 2010). This DIC source and control pathway has also been reported for estuarine environments (Raymond et al. 2000). Therefore, in the Everglades, respiration during the metabolism of organic matter is the primary source of DIC in surface waters, with organic matter, nutrient gradients and DOC concentrations controlling C turnover as indicated by the vary large *p*CO_2(aq)_ concentrations observed (Fig. 3).

Odum et al. (1979) hypothesized that in wetlands, plant productivity will be greatest when periodic short duration flooding provides subsidies of nutrients and water. Alternatively, prolonged flooding will cause physiological stress to the plants and limit nutrient subsidies thus limiting productivity. Using this subsidy-stress conceptual model, hydrology as indicated by hydroperiod is highly variable across the Everglades landscape (Appendix 2, Fig 2) combined with a strong gradient of available nutrients such as the WCA-2 eutrophication gradient (DeBusk et al. 2001; Julian et al. 2016). The eutrophication gradient itself is a combination of both “toxic” and “useable”-inputs as explained by the hypothetical performance curve of a perturbed ecosystem in this subsidy-stress concept. Useable inputs enhance productivity, alternatively, toxic inputs causes rapid declines in response to perturbation. Areas nearest the inflow experience stress from the availability of nutrients with impacts identified by significant shifts in species composition and biogeochemical processes (Reddy et al. 1993; Qualls and Richardson 2000) resulting in a significant decline in oligotrophic indicator species (i.e. sawgrass, calcareous periphyton) and suitable conditions for eutrophic indicator species (i.e. cattails). Meanwhile, further along the eutrophication gradient water availability is more dynamic, subsidized by a significant groundwater connection (Harvey et al. 2002) with nutrient concentrations gradually reach background concentrations and species composition and productivity return to oligotrophic conditions corroborated by observed DIC concentrations. Based on these observations the WCA-2 eutrophication gradient may exhibit subsidy-stress in both time and space and are determined by “chronic” nutrient availability and hydrologic variability. Likewise, WCA-1 and WCA-3 could also experience subsidy-stress conditions in light of variable hydrology and strong gradients of nutrient availability.

Nutrient availability influences the accretion of organic matter by stimulating net primary production and decomposition via microbial metabolism which in turn contributed to elevated DIC production, utilization and turnover (Reddy et al. 1993; Qualls and Richardson 2000; Fisher and Reddy 2010). Furthermore, the correlation of TN, specific conductance and DIC concentration (Table 4) could suggest evidence of a nutrient subsidy stimulated DIC production. Specific conductivity has been used as a tracer of surface water with higher conductivity water representing higher available nutrient canal water penetrating portions of the marsh while lower specific conductivity water represents interior marsh surface water with relatively low nutrient concentrations (Harwell et al. 2008; Surratt et al. 2008). The correlation of TN and DIC could be linked to the productivity of periphyton and blue-green algae which fix atmospheric nitrogen during metabolism. Finally, there is a hydrologic factor involved with DIC production as suggested by the wetland subsidy-stress model of Odum et al. (1979) in that WCA-1 is hydrologically isolated and WCA-3 is hydrologically fractured (i.e. compartmentalized), which in part regulates ecosystem level productivity, DIC production and carbon turnover (Table 4).

**Table 4.** Spearman correlation analysis results of monthly mean surface water *carbon variables* and various water quality parameters. Values are represented as spearman’s rho (ρ-value). Correlations that are not statistically significant are identified in bold text.

|  | Parameter | WCA-1 | WCA-2 | WCA-3 |
| --- | --- | --- | --- | --- |
| Dissolved Inorganic Carbon | Stage Elevation | **0.19 (0.14)** | **-0.25 (0.06)** | -0.49 (<0.01) |
|  | Water Temperature | **0.06 (0.63)** | -0.31 (<0.01) | -0.57 (<0.01) |
|  | Total Phosphorus | 0.39 (<0.01) | 0.34 (<0.05) | **-0.23 (0.07)** |
|  | Total Nitrogen | 0.62 (<0.01) | 0.65 (<0.01) | 0.59 (<0.01) |
|  | Specific Conductance | 0.74 (<0.01) | 0.67 (<0.01) | 0.65 (<0.01) |
| Dissolved Organic Carbon | Stage Elevation | **0.07 (0.61)** | -0.33 (<0.05) | -0.28 (<0.05) |
|  | Water Temperature | **-0.08 (0.52)** | **-0.14 (0.30)** | **-0.03 (0.79)** |
|  | Total Phosphorus | **0.25 (0.05)** | 0.38 (<0.01) | **0.009 (0.95)** |
|  | Total Nitrogen | 0.78 (<0.01) | 0.67 (<0.01) | 0.72 (<0.01) |
|  | Specific Conductance | 0.74 (<0.01) | 0.52 (<0.01) | 0.60 (<0.01) |
| Particulate Organic Carbon | Stage Elevation | **-0.23 (0.10)** | **-0.007 (0.96)** | **0.14 (0.28)** |
|  | Water Temperature | **-0.15 (0.26)** | **-0.12 (0.37)** | **-0.18 (0.18)** |
|  | Total Phosphorus | **0.08 (0.55)** | 0.40 (<0.01) | 0.37 (<0.01) |
|  | Total Nitrogen | **0.19 (0.16)** | **0.17 (0.23)** | **-0.11 (0.41)** |
|  | Specific Conductance | 0.30 (<0.05) | **0.04 (0.79)** | -0.32 (<0.05) |
| $p\text{CO}_{2,\text{aq}}$ | Stage Elevation | **-0.05 (0.20)** | **0.25 (0.06)** | **0.19 (0.14)** |
|  | Water Temperature | **0.25 (0.05)** | 0.35 (<0.01) | **0.21 (0.10)** |
|  | Total Phosphorus | 0.30 (<0.05) | 0.48 (<0.01) | **0.22 (0.09)** |
|  | Total Nitrogen | **0.22 (0.10)** | **0.08 (0.51)** | **0.09 (0.48)** |
|  | Specific Conductance | **0.12 (0.38)** | **0.02 (0.91)** | **-0.13 (0.31)** |

### Dissolved Organic Carbon and Particulate Organic Carbon

Dissolved organic C concentrations during this study were consistent with concentrations observed in other studies within the Everglades system (Qualls and Richardson 2003; Aiken et al. 2011; Lu et al. 2014). Wetlands are major sinks of C that sequester 0.003 – 2.2 kg C m^-2^ yr^-1^, with DOC typically representing <1% of the TOC in soil (i.e. bulk soil + porewater), but ~90% of the TOC in surface water (Kadlec and Wallace 2009; Kayranli et al. 2010). Particulate organic C concentrations observed during this study corroborate the consensus that DOC is the dominate form of organic C in the water column, typically an order of magnitude greater than POC concentrations (Fig. 4 and Table 3). Furthermore, suspended particulate concentrations in the interior portions of the EPA are relatively low (Julian et al. 2016), providing more evidence which suggests that most of the organic C is in the dissolved fraction (Lu et al. 2003).

The size of the DOC pool is orders of magnitude larger than the POC pool within the northern portion of the EPA. Nutrients can be a controlling factor in the production of organic C via aquatic plant and algae production, and decomposition. Surface water nutrient concentrations were positively correlated with both DOC and POC (Table 4), suggesting higher primary productivity in the water column and rates of organic C leaching and litterfall decomposition in areas of higher nutrients occurs within the Everglades system which correspond to result presented by previous studies (DeBusk and Reddy 1998; D’Angelo and Reddy 1999; Stern et al. 2007). Additionally, marsh TP concentrations in the WCAs have significantly decreased since WY1979 with WCA-2 having the largest period of record trend decrease (WY1979-WY2015; - 0.48 μg L^-1^ yr^-1^) in concentrations (Julian et al. 2016). The decrease in nutrient concentrations also corresponded with significant decreases in DOC, POC, DIC and *p*CO_2_ (this study; Table 2) with WCA-2 observing the largest concurrent decreases across the EPA. Availability of nutrients and electron acceptors plays an important role in wetland C cycling with available nutrients limiting the growth of microbial fauna (Drake et al. 1996; D’Angelo and Reddy 1999; Wright and Reddy 2001).

While nutrients play an important role in DIC production and C turnover, hydrologic conditions also play a significant role in the cycling and production of organic C within wetlands. Dissolved organic C was negatively correlated with stage elevation, suggesting a possible dilution of DOC within marshes (Table 4). (Lu et al. 2003) observed a similar correlation with DOC in the southern portion of ENP, suggesting that rainfall diluted DOC concentrations as water levels increases along flow transects within the Everglades system. This process has also been demonstrated in other forested and wetland ecosystems (Fraser et al. 2001; Kawasaki et al. 2005; Ågren et al. 2007). Some studies have attributed this decreases in DOC being caused by flushing and export of DOC from the system as seen in forested ecosystems (Inamdar et al. 2004). Additionally, (Lu et al. 2003) observed that canal inflow water is a major source of dissolved organic matter in wetlands, the majority of which is DOC. As the canal surface water flows penetrated through marsh DOC is leached from periphyton, vegetation, senescent plant material, detritus and soil which increased the marsh DOC concentrations. The strong correlation between DOC and specific conductance provides evidence towards this DOC dynamic (Table 4). Using radio-isotopes, (Stern et al. 2007) determined that DOC turnover rates are typically longer than the wetland water residence time, suggesting that most of the DOC will persist in the water column as the water flows. Therefore, the temporal and spatial variation in marsh DOC concentrations and to some extent POC concentrations are driven in part by the source of water and factors contributing to the flux of DOC from ecosystem components to the overlying water column.

Particulate OC has been used to estimate carbon burial in oceanic, estuarine, lake and stream ecosystems (Schindler et al. 1997; Barth et al. 1998; Robertson et al. 1999; Allison et al. 2007). Rivers and streams are net transporters of POC to lake and estuarine ecosystems. In deep lakes and marine environments POC is an important source of C and quickly mineralized.

Accumulation rates can vary several orders of magnitude across systems ranging from 0.9 g C m^-2^ yr^-1^ in riverine tributaries (Kao and Liu 1996) to 378 g C m^-2^ yr^-1^ in oceanic systems (Suess 1980). As discussed above, DOC is the dominate fraction in wetland ecosystems (Wetzel 1984) however C accumulation rates in the Everglades is on par with most riverine ecosystems at an average rate of 205 g C m^-2^ yr^-1^ (range: 86 – 424 g C m^-2^ yr^-1^ (Reddy et al. 1993). Although in river ecosystems, POC concentrations and C accumulation rates can be relatively high due to allochthonous inputs of C while POC concentrations within the Everglades are relatively low and C accumulation rates are relatively moderate suggesting dominate authocthonous production of C.

### Dissolved Inorganic Carbon: Dissolved Organic Carbon Ratio

Each WCA experiences different hydrologic conditions. Some studies have attributed differences in DOC:DIC ratios as indicators of hydrologic influence on dissolved C flux dynamics in stream and river ecosystems (Elder et al. 2000; Palmer et al. 2001; Kawasaki et al. 2005). The data suggest that the hydrologic driver is also present in WCAs. Despite similar ecological features (i.e. ridge and slough), C loads and cycling differ between WCAs (Fig. 4). Furthermore, DOC:DIC ratios significantly differ between areas with the DOC:DIC ratio for WCA-1 deviating from WCA-2 and WCA-3 indicating greater DOC concentrations relative to DIC within WCA-1. Meanwhile, DOC:DIC ratios within WCA-2 and WCA-3 indicate a more balanced dissolved C flux (Fig. 5) but differ slightly presumable due to differences in water quality conditions (Table 3). The difference of WCA-1 to the other WCAs is primarily attributed to differences in hydrologic condition with interior portions of WCA-1 receiving very little surface water flow from canals but rather rainfall, which has low alkalinity concentrations and relatively low pH (Julian et al. 2016). Meanwhile, WCA-2 and WCA-3 are driven largely by surface water and groundwater flow (Harvey et al. 2004; Harvey and McCormick 2009).

In lake ecosystems decomposition, mineralization and sedimentation processes would reduce DOC and increase DIC, which would overall decreases the DOC:DIC ratios. Meanwhile, surface water and groundwater inputs into lakes can potentially increase DOC:DIC ratios depending on the source and magnitude of flow (Elder et al. 2000). Lake waters are commonly supersaturated with respect to CO_2_ as a result of CO_2_ generated from the decomposition of organic matter imported from upstream, resulting in a net flux of C to the atmosphere (Cole et al. 1994, 2001). A similar process occurs in wetland, wetlands especially the Everglades receive copious quantities of allochthonous C and produce large quantities of autochthonous C from biomass turnover therefore allowing the water column to become supersaturated with CO_2_ resulting in large CO_2_ fluxes (Table 3).

### CO_2(aq)_ versus CO_2(atm)_

Over the past century, the mean global CO_2(atm)_ has risen from approximately 280 μatm to over 368 μatm (Keeling and Whorf 1994; Baldocchi et al. 2001). Similarly, atmospheric CO_2_ concentrations increases across the study period at the Key Biscayne monitoring location. While this location is ~60 kilometers from the study location (center of WCA-3 to Key Biscayne), atmospheric CO_2_ concentrations observed at Key Biscayne are comparable to data collected from locations within ENP (J.G. Barr, *Unpublished data* and G. Starr, *Unpublished data*) at three separate monitoring locations with a more limited period of record. In contrast, annual mean *p*CO_2(aq)_ concentrations significantly declined within the WCAs (Table 2). Even though *p*CO_2(aq)_ is orders of magnitude greater than that of the *p*CO_2(atm)_ this diverging relationship is unexpected and more work is needed to explore to explain this phenomenon.

Wetlands release large quantities of C as CO_2_ (Table 3) and methane due to anaerobic conditions and the decomposition of organic matter. In wetlands, C sequestration typically outpaces C release making wetland soils the world’s largest C sinks (Kayranli et al. 2010). Here, the difference between *CO_2(aq)_ versus CO_2(atm)_* indicates increasing flux to the atmosphere. This does not mean that these ecosystems are net sources as this can be compensated by a particulate flux from emergent vegetation or by allochthonous carbon. However, the temporal pattern suggests long-term changes.

Wetland C reserves are at risk of becoming atmospheric C sources due to changes in climate and hydrologic conditions (Gorham 1991). Climate change in the form of increased temperatures and altered precipitation patterns (Kundzewicz et al. 2008; Whitehead et al. 2009) can potentially decrease wetlands ability to store C. Increased temperatures could stimulate organic matter decomposition and reduced rainfall and surface water flow into wetlands due to a drier climate could lead to compaction of organic soils allowing for rapid subsidence (Reddy et al. 2006; Kayranli et al. 2010). If the projected future climate is expected to be drier than past climates, WCA-1 could be the most affected by climate change due to its isolated hydrology, low DIC and high CO_2_ flux rates. Currently, it is not certain how climate change will influence C storage and other ecosystem processes of the Everglades and more direct research is needed to explore the effects of climate change and sea-level rise on C storage and processes in the Everglades ecosystem.

### Conclusion

Hydrologic condition, nutrient inputs, and nutrient cycling significantly influence the balance, speciation, and flux of C from wetland ecosystems consistent with the hypothesized subsidy-stress model proposed by (Odum et al. 1979) for wetland vegetation. Within the Everglades ecosystem the interplay between nutrient inputs and hydrologic condition exert a driving force on the balance between DIC and DOC production via the metabolism of organic matter. As Everglades restoration efforts progress and water quality continues to improve within the Everglades ecosystem the C cycle and associated C pools will also respond in kind. More specifically, impacted portions of the WCAs that have achieved a long-term TP concentration of 10 μg P L^-1^ recovering from impacted to unimpacted (Julian 2015) also exhibited significant declines in DIC concentrations. However, in light of expected climate change (i.e. altered precipitation, warmer air temperatures, etc.), the mechanisms of C pool control and production could change as a result of drier climates could significantly impact rainfall driven portions of the Everglades such as WCA-1. Perhaps future research can identify direct linkages of DOC, DIC and climate change trends as forecasts for the Everglades Ecosystem.

## Acknowledgements

We would like to thank the anonymous peer-reviewer(s) and editor(s) for their efforts and constructive review of this manuscript. Additionally, we would like to acknowledge all of the current and past South Florida Water Management District staff involved in the collection and laboratory analysis of the data used in this manuscript.

## Authors’ Contributions

PJ designed the study, performed data analysis including necessary calculations and statistics analysis, and wrote the manuscript. SG assisted in data analysis and provided editorial assistance. BG, ALW and TZO provided substantial editors assistance and review. All authors read and approved the final manuscript.

## References

Ågren A, Buffam I, Jansson M, Laudon H (2007) Importance of seasonality and small streams for the landscape regulation of dissolved organic carbon export. J Geophys Res Biogeosciences 112:G03003. 10.1029/2006JG000381

Aiken GR, Hsu-Kim H, Ryan JN (2011) Influence of Dissolved Organic Matter on the Environmental Fate of Metals, Nanoparticles, and Colloids. Environ Sci Technol 45:3196–3201. 10.1021/esl03992s

Allison MA, Bianchi TS, McKee BA, Sampere TP (2007) Carbon burial on river-dominated continental shelves: Impact of historical changes in sediment loading adjacent to the Mississippi River. Geophys Res Lett 34:L01606. 10.1029/2006GL028362

Baldocchi D, Falge E, Gu L, et al (2001) FLUXNET: A New Tool to Study the Temporal and Spatial Variability of Ecosystem–Scale Carbon Dioxide, Water Vapor, and Energy Flux Densities. Bull Am Meteorol Soc 82:2415–2434. 10.1175/1520-0477(2001)082<2415:FANTTS>2.3.CO;2

Bancroft GT, Hoffman W, Sawicki RJ, Ogden JC (1992) The Importance of the Water Conservation Areas in the Everglades to the Endangered Wood Stork (Mycteria americana). Conserv Biol 6:392–398. 10.1046/j.1523-1739.1992.06030392.x

Barth JAC, Veizer J, Mayer B (1998) Origin of particulate organic carbon in the upper St. Lawrence: isotopic constraints. Earth Planet Sci Lett 162:111–121. 10.1016/S0012-821X(98)00160-5

Billett MF, Moore TR (2008) Supersaturation and evasion of CO2 and CH4 in surface waters at Mer Bleue peatland, Canada. Hydrol Process 22:2044–2054. 10.1002/hyp.6805

Bossio DA, Fleck JA, Scow KM, Fujii R (2006) Alteration of soil microbial communities and water quality in restored wetlands. Soil Biol Biochem 38:1223–1233. 10.1016/j.soilbio.2005.09.027

Brandt LA, Portier KM, Kitchens WM (2000) Patterns of change in tree islands in Arthur R. Marshall Loxahatchee National Wildlife Refuge from 1950 to 1991. Wetlands 20:1–14. 10.1672/0277506 5212(2000)020[0001:POCITI]2,O.CO;2

Bronaugh DB, Werner A (2013) Zhang + Yue-Pilon trends package. CRAN R-Project

Bruland GL, Grunwald S, Osborne TZ, et al (2006) Spatial Distribution of Soil Properties in Water Conservation Area 3 of the Everglades. Soil Sci Soc Am J 70:1662. 10.2136/sssaj2005.0134

Carpenter SR, Pace ML (1997) Dystrophy and Eutrophy in Lake Ecosystems: Implications of Fluctuating Inputs. Oikos 78:3. 10.2307/3545794

Cole JJ, Caraco NF, Kling GW, Kratz TK (1994) Carbon Dioxide Supersaturation in the Surface Waters of Lakes. Science 265:1568–1570.

Cole JJ, Cole JJ, Caraco NF, Caraco NF (2001) Carbon in catchments: connecting terrestrial carbon losses with aquatic metabolism. Mar Freshw Res 52:101–110.

Cole JJ, Pace ML, Carpenter SR, Kitchell JF (2000) Persistence of net heterotrophy in lakes during nutrient addition and food web manipulations. Limnol Oceanogr 45:1718–1730. 10.4319/IO.2000.45.8.1718

Corstanje R, Reddy KR, Prenger JP, et al (2007) Soil microbial eco-physiological response to nutrient benrichment in a sub-tropical wetland. Ecol Indic 7:277–289. 10.1016/j.ecolind.2006.02.002

D’Angelo EM, Reddy KR (1999) Regulators of heterotrophic microbial potentials in wetland soils. Soil Biol Biochem 31:815–830. 10.1016/S0038-0717(98)00181-3

DeBusk WF, Newman S, Reddy KR (2001) Spatio-temporal patterns of soil phosphorus enrichment in Everglades Water Conservation Area 2A. J Environ Qual 30:1438–1446.

DeBusk WF, Reddy KR (1998) Turnover of detrital organic carbon in a nutrient-impacted Everglades bmarsh. Soil Sci Soc Am J 62:1460–1468.

Dinno A (2015) Dunn’s test of multiple comparisons using rank sums. CRAN R-Project

Drake HL, Aumen NG, Kuhner C, et al (1996) Anaerobic Microflora of Everglades Sediments: Effects of Nutrients on Population Profiles and Activities. Appl Environ Microbiol 62:486–493.

Duarte CM, Cebrian J (1996) The fate of marine autotrophic production. Limnol Oceanogr 41:1758–531–1766.

Elder JF, Rybicki NB, Carter V, Weintraub V (2000) Sources and yields of dissolved carbon in northern Wisconsin stream catchments with differing amounts of peatland. Wetlands 20:113–125. 10.1672/0277-5212(2000)020[0113:SAYODC]2,O.CO;2

Elser JJ, Bracken MES, Cleland EE, et al (2007) Global analysis of nitrogen and phosphorus limitation of primary producers in freshwater, marine and terrestrial ecosystems. Ecol Lett 10:1135–1142. 10.1111/j.1461-0248.2007.01113.x

Findlay SEG, Dye S, Kuehn KA (2002) Microbial growth and nitrogen retention in litter of Phragmites australis compared to Typha angustifolia. Wetlands 22:616–625. 10.1672/0277540 5212(2002)022[0616:MGANRI]2,O.CO;2

Finlay JC (2003) Controls of streamwater dissolved inorganic carbon dynamics in a forested watershed. Biogeochemistry 62:231–252. 10.1023/A:1021183023963

Fisher MM, Reddy KR (2010) Estimating the Stability of Organic Phosphorus in Wetland Soils. Soil Sci Soc Am J 74:1398. 10.2136/sssaj2009.0268

Florida Administrative Code (2008) Chapter 62-160 Quality Assurance.

Fraser CJD, Roulet NT, Moore TR (2001) Hydrology and dissolved organic carbon biogeochemistry in an ombrotrophic bog. Hydrol Process 15:3151–3166. 10.1002/hyp.322

Freeman C, Fenner N, Ostle NJ, et al (2004) Export of dissolved organic carbon from peatlands under elevated carbon dioxide levels. Nature 430:195–198. 10.1038/nature02707

Gleason PJ, Cohen AD, Smith WG, et al (1974) The environmental significance of Holocene sediments from the Everglades and saline tidal plain. In: Environments of south Florida: Present and past. Miami Geological Society: Coral Gables, FL, pp 287–341

Gorham E (1991) Northern peatlands: role in the carbon cycle and probable responses to climatic warming. Ecol Appl 1:182–195.

Gu B, Miao S, Edelstein C, Dreschel T (2008) Effects of a prescribed fire on dissolved inorganic carbon dynamics in a nutrient-enriched Everglades wetland. Fundam Appl Limnol Arch Für Hydrobiol 171:263–272. 10.1127/1863-9135/2008/0171-0263

Gu B, Schelske CL, Coveney MF (2011) Low carbon dioxide partial pressure in a productive subtropical lake. Aquat Sci 73:317–330. 10.1007/s00027-010-0179-γ

Hagerthey SE, Cole JJ, Kilbane D (2010) Aquatic metabolism in the Everglades: Dominance of water column heterotrophy. Limnol Oceanogr 55:653–666.

Hanson PC, Bade DL, Carpenter SR, Kratz TK (2003) Lake metabolism: Relationships with dissolved organic carbon and phosphorus. Limnol Oceanogr 48:1112–1119. 10.4319/IO.2003.48.3.1112

Harvey JW, Krupa SL, Gefvery C, et al (2002) Interactions between surface water and groundwater effects on mercury transport in the noeth-central Everglades. United States Geological Survey, Reston, VA

Harvey JW, Krupa SL, Krest JM (2004) Ground Water Recharge and Discharge in the Central Everglades. Ground Water 42:1090–1102. 10.1111/j.1745-6584.2004.tb02646.x

Harvey JW, McCormick PV (2009) Groundwater’s significance to changing hydrology, water chemistry, and biological communities of a floodplain ecosystem, Everglades, South Florida, USA. Hydrogeol J 17:185–201. 10.1007/sl0040-008-0379-x

Harwell MC, Surratt DD, Barone DM, Aumen NG (2008) Conductivity as a tracer of agricultural and urban runoff to delineate water quality impacts in the northern Everglades. Environ Monit Assess 147:445–462. 10.1007/sl0661-007-0131-3

Hessen DO, Ågren GI, Anderson TR, et al (2004) Carbon sequestration in ecosystems: the role of stoichiometry. Ecology 85:1179–1192. 10.1890/02-0251

Hopkinson CS, Cai W-J, Hu X (2012) Carbon sequestration in wetland dominated coastal systems — a global sink of rapidly diminishing magnitude. Curr Opin Environ Sustain 4:186–194. 10.1016/j.cosust.2012.03.005

Inamdar SP, Christopher SF, Mitchell MJ (2004) Export mechanisms for dissolved organic carbon and nitrate during summer storm events in a glaciated forested catchment in New York, USA. Hydrol Process 18:2651–2661. 10.1002/hyp.5572

Julian P (2015) Appendix 3A-6: Water Year 2010-2014 annual total phosphorus criteria compliance assessment. In: 2015 South Florida Environmental Report. South Florida Water Management District, West Palm Beach, FL,

Julian P, Payne GG, Xue SK (2016) Chapter 3A: Water Quality in the Everglades Protection Areas. In: 2016 South Florida Environmental Report. South Florida Water Management District, West Palm Beach, FL,

Kadlec RH, Wallace SD (2009) Treatment wetlands. CRC Press, Boca Raton, FL

Kao S-J, Liu K-K (1996) Particulate organic carbon export from a subtropical mountainous river (Lanyang Hsi) in Taiwan. Limnol Oceanogr 41:1749–1757. 10.4319/lo.1996.41.8.1749

Kawasaki M, Ohte N, Katsuyama M (2005) Biogeochemical and hydrological controls on carbon export from a forested catchment in central Japan. Ecol Res 20:347–358. 10.1007/sl1284-0055950050-0

Kayranli B, Scholz M, Mustafa A, Hedmark Å (2010) Carbon Storage and Fluxes within Freshwater Wetlands: a Critical Review. Wetlands 30:111–124. 10.1007/sl3157-009-0003-4

Keeling CD, Whorf TP (1994) Atmospheric CO 2 records from sites in the SIO air sampling network, Trends’ 93: A Compendium of Data on Global Change TA Boden, et al. Rep. Oak Ridge National Laboratory, Oak Ridge, Tennessee

Kling GW, Kipphut GW, Miller MC (1991) Arctic Lakes and Streams as Gas Conduits to the Atmosphere: Implications for Tundra Carbon Budgets. Science 251:298.

Kling GW, Kipphut GW, Miller MC (1992) The flux of CO2 and CH4 from lakes and rivers in arctic Alaska. Hydrobiologia 240:23–36. 10.1007/BF00013449

Kundzewicz ZW, Mata LJ, Arnell NW, et al (2008) The implications of projected climate change for freshwater resources and their management. Hydrol Sci J 53:3–10. 10.1623/hysj.53.1.3

Light SS, Dineen JW (1994) Water control in the everglades: a historical perspective. In: Davis S, Ogden J (eds) Everglades: The ecosystem and its restoration. St. Lucie Press, Delray Beach, FL, pp 47–84

Lu H, Yang L, Shabbir S, Wu Y (2014) The adsorption process during inorganic phosphorus removal by cultured periphyton. Environ Sci Pollut Res 21:8782–8791. 10.1007/sll356-014-2813-z

Lu XQ, Maie N, Hanna JV, et al (2003) Molecular characterization of dissolved organic matter in freshwater wetlands of the Florida Everglades. Water Res 37:2599–2606. 10.1016/S0043-1354(03)00081-2

Newman S, Hagerthey SE (2011) Water Conservation Area 1: A Case Study of Hydrology, Nutrient, and Mineral Influences on Biogeochemical Processes. Crit Rev Environ Sci Technol 41:702–722. 10.1080/10643389.2010.530910

Odum EP, Finn JT, Franz EH (1979) Perturbation Theory and the Subsidy-Stress Gradient. BioScience 29:349–352. 10.2307/1307690

Odum HT (1956) Primary production in flowing waters. Limnol Ocean 1:102–117.

Palmer SM, Hope D, Billett MF, et al (2001) Sources of organic and inorganic carbon in a headwater stream: Evidence from carbon isotope studies. Biogeochemistry 52:321–338. 10.1023/A:1006447706565

Qualls RG, Richardson CJ (2000) Phosphorus enrichment affects litter decomposition, immobilization, and soil microbial phosphorus in wetland mesocosms. Soil Sci Soc Am J 64:799–808.

Qualls RG, Richardson CJ (2003) Factors controlling concentration, export, and decomposition of dissolved organic nutrients in the Everglades of Florida. Biogeochemistry 62:197–229. 10.1023/A:1021150503664

Raven J, Beardall J, Griffiths H (1982) Inorganic C-sources for Lemanea, Cladophora and Ranunculus in a fast-flowing stream: Measurements of gas exchange and of carbon isotope ratio and their ecological implications. Oecologia 53:68–78. 10.1007/BF00377138

Raymond PA, Bauer JE, Cole JJ (2000) Atmospheric C02 evasion, dissolved inorganic carbon production, and net heterotrophy in the York River estuary. Limnol Oceanogr 45:1707–1717. 10.4319/IO.2000.45.8.1707

Reddy KR, DeLaune RD, DeBusk WF, Koch MS (1993) Long-term nutrient accumulation rates in the Everglades. Soil Sci Soc Am J 57:1147–1155.

Reddy KR, Osborne TZ, Inglett KS, Corstanje R (2006) Influence of water levels on subsidence of organic soils in the upper St. Johns River Basin. Saint Johns River Water Management District, Palatka, FL, USA

Reddy KR, Wang Y, DeBusk WF, et al (1998) Forms of soil phosphorus in selected hydrologic units of the Florida Everglades. Soil Sci Soc Am J 62:1134–1147.

Richey JE, Melack JM, Aufdenkampe AK, et al (2002) Outgassing from Amazonian rivers and wetlands as a large tropical source of atmospheric CO2. Nature 416:617–620. 10.1038/416617a

Robertson Al, Bunn SE, Boon PI, Walker KF (1999) Sources, sinks and transformations of organic carbon in Australian floodplain rivers. Mar Freshw Res 50:813. 10.1071/MF99112

Sand-Jensen K, Frost-Christensen H (1998) Photosynthesis of amphibious and obligately submerged plants in CO2-rich lowland streams. Oecologia 117:31–39. 10.1007/s004420050628

Scheidt D, Kalla PI (2007) Everglades ecosystem assessment: water management and quality, eutrophication, mercury contamination, soil and habitat: monitoring for adapyive management: a R-EMAP status report. United States Environmental Protection Agency, Athens, GA

Schindler DW, Curtis PJ, Bayley SE, et al (1997) Climate-Induced Changes in the Dissolved Organic Carbon Budgets of Boreal Lakes. Biogeochemistry 36:9–28.

Stern J, Wang Y, Gu B, Newman J (2007) Distribution and turnover of carbon in natural and constructed wetlands in the Florida Everglades. Appl Geochem 22:1936–1948. 10.1016/j.apgeochem.2007.04.007

Suess E (1980) Particulate organic carbon flux in the oceans—surface. Nature 288:261.

Surratt DD, Waldon MG, Harwell MC, Aumen NG (2008) Time-series and spatial tracking of polluted canal water intrusion into wetlands of a National Wildlife Refuge in Florida, USA. Wetlands 28:176–183. 10.1672/07-74.1

Telmer K, Veizer J (1999) Carbon fluxes, pC02 and substrate weathering in a large northern river basin, Canada: carbon isotope perspectives. Chem Geol 159:61–86. 10.1016/S0009-2541(99)00034-0

Updegraff K, Pastor J, Bridgham SD, Johnston CA (1995) Environmental and Substrate Controls over Carbon and Nitrogen Mineralization in Northern Wetlands. Ecol Appl 5:151–163. 10.2307/1942060

Waiser M, Robarts R (2004) Net heterotrophy in productive prairie wetlands with high DOC concentrations. Aquat Microb Ecol 34:279–290. 10.3354/ame034279

Wetzel RG (1992) Gradient-dominated ecosystems: sources and regulatory functions of dissolved organic matter in freshwater ecosystems. In: Salonen K, Kairesalo T, Jones RI (eds) Dissolved Organic Matter in Lacustrine Ecosystems. Springer Netherlands, pp 181–198

Wetzel RG (1984) Detrital Dissolved and Particulate Organic Carbon Functions in Aquatic Ecosystems. Bull Mar Sci 35:503–509.

Wetzel RG, Likens GE (2000) The Inorganic Carbon Complex: Alkalinity, Acidity, CO2, pH, Total Inorganic Carbon, Hardness, Aluminum. In: Limnological Analyses. Springer New York, pp 113–135

Whitehead PG, Wilby RL, Battarbee RW, et al (2009) A review of the potential impacts of climate change on surface water quality. Hydrol Sci J 54:101–123. 10.1623/hysj.54.1.101

Wright AL, Reddy KR (2001) Heterotrophic Microbial Activity in Northern Everglades Wetland Soils. Soil Sci Soc Am J 65:1856. 10.2136/sssaj2001.1856

